# Hydrological balance of a subalpine forest and the effects of fog presence and forest age

**DOI:** 10.64898/2026.05.07.723430

**Authors:** Glenda Garcia-Santos, Nikolaus Obojes, Leonardo Montagnani

## Abstract

Subalpine forests in the Alps are fragile ecosystems that play a crucial role in regional water resources and the local climate. These ecosystems are ecologically significant due to their unique biodiversity and vulnerability to climate change. While several components of the hydrological balance have been studied, the interplay between catchment-scale processes and plot-scale drivers such as fog presence and forest age remains insufficiently understood.

To address this, we investigated the hydrological balance of a subalpine coniferous forest catchment at the Renon site in the Italian Alps, integrating observations across spatial scales. The study area includes a mosaic of mature and younger regrowth forest, where both interannual and seasonal variability in precipitation and fog presence are pronounced. At the catchment scale, we quantified above-canopy precipitation, evapotranspiration (ET, measured via eddy covariance at the ICOS tower), stream discharge, and soil moisture dynamics.

Within the catchment, we characterised water partitioning using sap flow sensors for tree transpiration, throughfall and stemflow collectors with rain gauges above and below the canopy and epiphyte sampling. Mixed fog-rain events frequently coincided with higher throughfall. However, these changes had a minor effect on soil water storage and catchment discharge in the annual water balance, which was nearly closed. At the plot scale, our results show that tree transpiration was higher in the younger forest structure, while canopy interception is a dominant process in water partitioning in the older forest structure, where lichen abundance likely enhances interception. This study highlights the importance of multi-scale monitoring in temperate mountain forests, where forest age influences water partitioning, and fog presence, though not directly quantified, can still contribute to reducing evaporative processes. Such contributions may gain importance under changing climate conditions, albeit less prominently than in tropical or subtropical cloud forests.

## 1. Introduction

Mountain forests, particularly those in the European Alps, are critical regulators of hydrological processes and regional climate systems. They provide essential ecosystem services, including water retention, runoff moderation, and thermal buffering, which are increasingly important under ongoing and projected climate change (Messerli et al., 2004; Beniston, 2005; Gobiet et al., 2014). The Alps function as natural “water towers” for surrounding lowlands, storing and releasing water through precipitation interception, evapotranspiration, and soil-water interactions (Viviroli et al., 2007; Beniston et al., 2011). However, climate-induced warming, changes in precipitation regimes, and shifts in vegetation dynamics are expected to profoundly affect the water balance and ecosystem functioning in these high-elevation environments (Grabherr et al., 2010; Gobiet et al., 2014).

Subalpine forests, dominated by coniferous species such as *Picea abies*, occupy the elevation zone just below the alpine treeline. Their dense canopies play a pivotal role in modulating water fluxes by intercepting rainfall, facilitating ET, and altering the partitioning between runoff and infiltration (Bonan, 2008; Bohn et al., 2000). Structural attributes like canopy height, leaf area index (LAI), and crown complexity vary with forest age and species composition, influencing interception capacity and transpiration rates. These effects are particularly relevant in the eastern Alps, where precipitation patterns are diverse, and forest cover is extensive, such as in South Tyrol, where forests occupy nearly 50% of the land area (Hilpold et al., 2018). Understanding how different forest structures influence hydrological processes is essential for predicting ecohydrological responses to land-use and climate change. A major challenge in this context is linking plot-scale variability in forest characteristics to catchment-scale water balance, as the spatial heterogeneity of forest ecosystems introduces complexities in measuring and modeling water fluxes. Many existing studies focus either on tree-scale measurements (e.g., sap flow or interception) or on integrated catchment-scale assessments, often lacking the resolution to connect these two levels meaningfully (Savenije, 2004; van Dijk et al., 2015; Nelson et al., 2020). This gap is particularly pronounced in temperate coniferous forests, where dense canopies and stratified understories complicate direct measurement of individual fluxes and their interactions.

Fog, though rarely quantified directly, may play a modifying role in hydrological inputs in these environments. In tropical and subtropical cloud forests, fog has been shown to contribute significantly to water availability, with throughfall and canopy deposition increasing on foggy days (García-Santos, 2007; Eugster et al., 2006). In temperate mountain forests, fog events often coincide with rainfall and are thus harder to separate and quantify. Still, their occurrence may enhance throughfall and reduce evaporative losses by increasing canopy humidity (Wagner and Petitta, 2015). While fog-only deposition is likely minor in the Alps, the presence of fog during precipitation events, so-called “mixed” conditions, may indirectly alter partitioning processes and evaporative dynamics.

In addition, the role of epiphytic organisms such as lichens and bryophytes deserves attention. These poikilohydric organisms absorb water passively from the atmosphere and are common in old-growth coniferous forests. By increasing the canopy water storage capacity and modifying drying rates, epiphytes may influence interception losses and contribute to latent heat fluxes (Pypker et al., 2006; Porada et al., 2018; Sillett and Van Pelt, 2007). Although their contribution to water fluxes remains poorly quantified, they likely represent a structurally mediated pathway through which forest age and fog presence jointly influence ecohydrological behaviour.

Forest age itself affects not only canopy structure but also flux partitioning. Younger forests tend to have high tree density and more homogeneous crowns, potentially leading to higher transpiration per unit area, whereas older forests may have more complex structures, greater epiphytic cover, and enhanced interception (Pypker et al., 2006). The implications of these differences for water balance remain insufficiently understood in the Alpine context.

Recent advances in ecohydrological instrumentation allow for detailed partitioning of forest water fluxes. Eddy covariance systems provide continuous ET measurements at the ecosystem scale, while sap flow sensors, throughfall and stemflow collectors, and soil moisture probes enable high-resolution monitoring at the plot level (Rebmann et al., 2018; Wieser et al., 2018). When deployed together in a nested monitoring design, these tools make it possible to link plot-level processes to catchment-scale hydrological functioning - a critical step for understanding the role of forest structure and microclimatic factors like fog.

In this study, we investigate the hydrological balance of a subalpine coniferous forest catchment in the eastern Alps, integrating flux measurements at both the catchment and plot scale. Our study site features a mosaic of forest patches of different ages and structural characteristics, allowing us to explore how forest age and canopy complexity influence water partitioning. Specifically, we address the following research questions:

1. How is the annual hydrological balance of a subalpine coniferous forest catchment partitioned among precipitation, evapotranspiration, discharge, and soil water storage change?
2. What are the magnitude and seasonal dynamics of evapotranspiration and tree transpiration in the young and old forest patches?
3. How is precipitation partitioned at the plot (canopy) scale, and how do forest age, epiphytic lichens, and fog occurrence during precipitation events modify throughfall and interception?

By combining long-term catchment monitoring with detailed plot-scale instrumentation, we aim to assess how structural forest variability and episodic fog presence contribute to precipitation partitioning and water balance closure in temperate coniferous mountain forests. This integrated, multi-scale approach contributes to a better understanding of forest ecohydrology under current and future climate conditions, and supports improved water management and forest conservation strategies.

## 2. Materials and methods

### 2.1 Study area

#### 2.1.1 Catchment description

The catchment is at the Renon-Grünwand site, South Tyrol, in the Italian Alps. It has an area of 0.44 km², at an elevation ranging from 1165 to 1918 m asl. The soil has developed above a layer of glacial till, with a depth of approximately 1 m, placed on top of a porphyry bedrock. The soil was classified as Haplic Podzol according to the FAO soil taxonomy (FAO, 1998) and, on average, consisted of 49% sand, 39% silt, and 12% clay. The catchment is characterised by a forest of natural origin and is managed for wood production. The traditional harvest method creates small gaps, approximately 50 m wide, and involves the thinning of surrounding trees. The result is a heterogeneous vegetation structure, with almost even-aged groups forming an uneven-aged structure with grassed forest gaps at a larger scale, as a result of the irregular cuttings of the several private properties present in the area.

#### 2.1.2 Plot

We selected the 0.9 ha area where the FluxNet research station named IT-Ren is located as a representative plot to study evapotranspiration, variation in soil water content, and the effects of fog presence. It is centred at (1735 m a.s.l., 46°35′11′′N, 11°26′00′′E, Montagnani et al., 2009), belonging to the ICOS infrastructure. In this area, we also considered the effects of fog presence, forest age and epiphytes. The diameters and heights of the trees in the research area have been measured every ten years since 1990, while the diameter of a subset of trees is measured annually by manual dendrometers (UMS, München, Germany). The most recent inventory (tree height, size, and position) was performed in 2020 (Badraghi et al., 2021) using a laser technique involving a UAV-mounted laser scanner flight with the TruPulse sensor (TruPulse 360 B laser range-finder; Laser Tech, Centennial CO, USA) (Figure 1).

**Figure 1.**
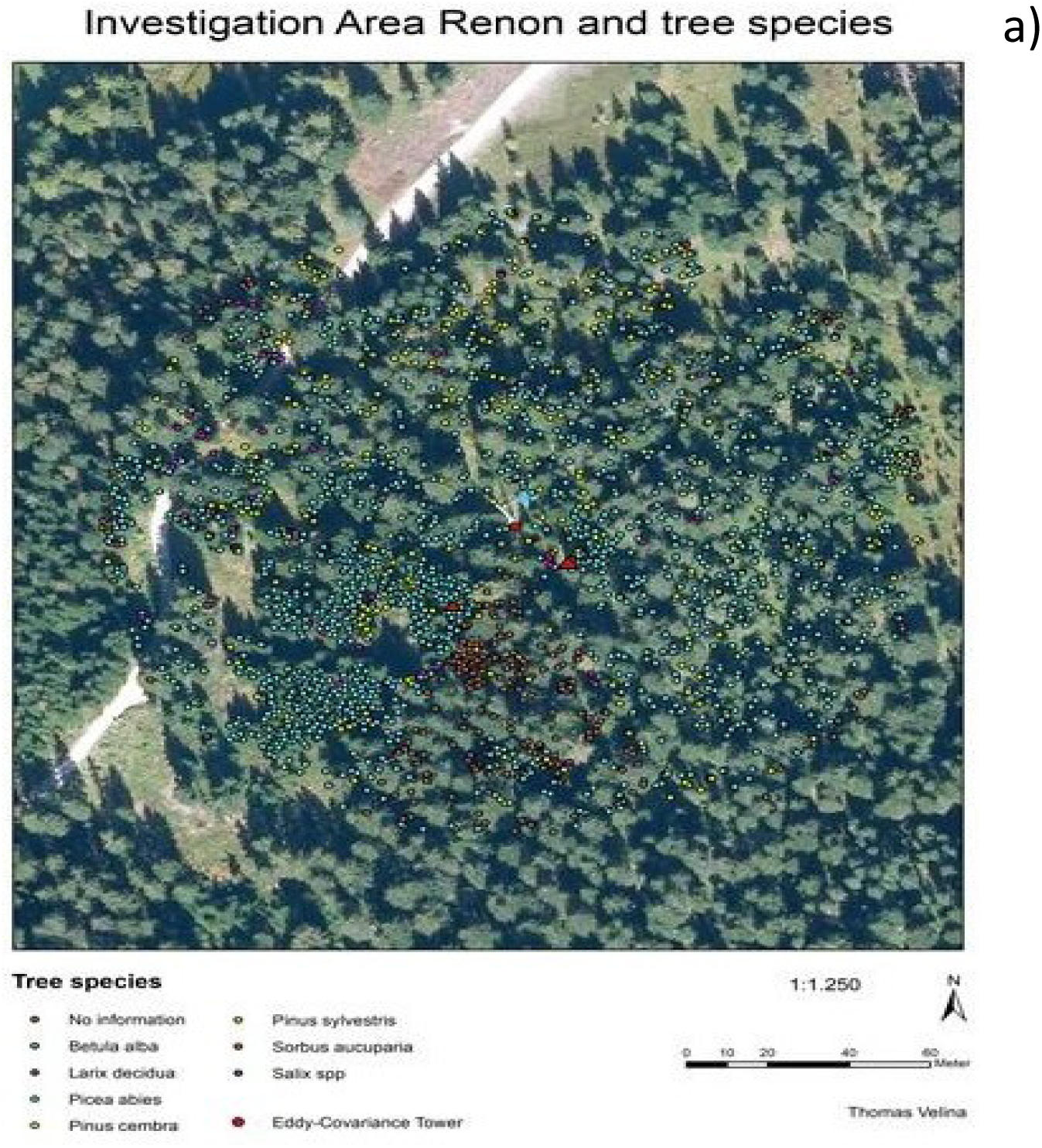
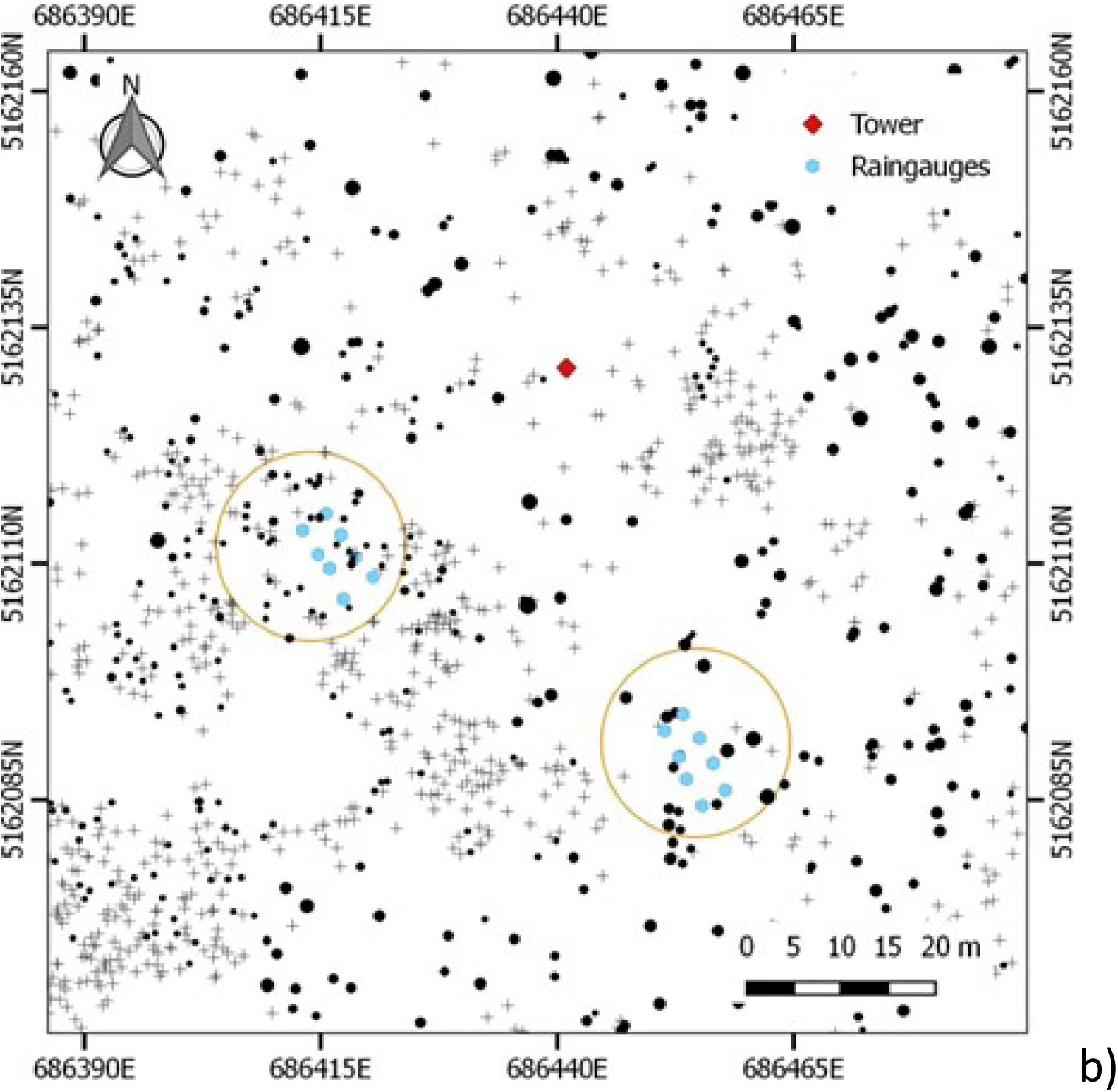
(a), Aerial image of the studied plot. (b), map of the trees, where we can see the sub-plots (red circles) selected as representative of young and old-growth patches of the forest. In the centre of the image, the eddy covariance tower is a red diamond. Throughfall measurements were conducted in both plots (blue points in the right figure), the young forest patch in the left and the old forest patch on the right.

The site is characterised by a large group of dominant spruce trees (*Picea abies (L) Karst.*) and, to a lesser extent, Swiss stone pine (*Pinus cembra* L.), with a small representation of European larch (*Larix europea* L.) trees (details in Table 1). Two groups can be differentiated, one with an age of approximately 200 years, and a second group of young, about 30-year-old trees. In both forest patches, parts of the living crown frequently reached the ground, irrespective of age. Local spruce and Swiss stone pine were characterised by an almost columnar shape (Marcolla et al., 2005). This shape and the high leaf area index (LAI, average 4.8) as measured by hemispherical images) created peculiar microclimatic conditions within the crown. The understory consisted mainly of blueberry (*Vaccinium myrtillus* L) and, in the grassed forest gaps, wavy hairgrass (*Deschampsia flexuosa*) was the dominant species.

**Table 1.** Indication of the spatial scales considered and the measured variables.

| Scale | Measured variables |
| --- | --- |
| Tree | Sap flow, epiphytes presence |
| Forest patch | Throughfall, stemflow, crown width, leaf area index |
| Plot | Forest inventory, soil water content, fog collection and multiannual fog occurrence statistics |
| Footprint | Evapotranspiration |
| Water basin | Water discharge |
| Landscape | Statistics about fog occurrence during the measurement year |

### 2.2 Hydrological monitoring concept

This study applies a nested observational design to integrate hydrological measurements at both catchment and plot scales. At the catchment scale, we monitored the water balance using measurements of precipitation, evapotranspiration (via eddy covariance), soil water content, and stream discharge during the whole of 2019. These variables were recorded continuously (Table 2) and used to estimate net water inputs, losses, and storage changes over the entire basin. Within the catchment, we selected two forest patches of contrasting age (approximately 200 years and 30 years, respectively) (Fig. 1) to capture plot-scale variability in forest structure and associated hydrological processes from 30.05.2019 until 07.11.2019. The contribution of epiphytic lichens to canopy water storage was also assessed. This design facilitates, on the one hand, the analysis of how variations in forest structure at the plot scale are reflected in the components of the catchment-scale water balance and, on the other hand, provides insights into the partitioning of evapotranspiration components and their implications for catchment-scale water balance closure.

**Table 2:**
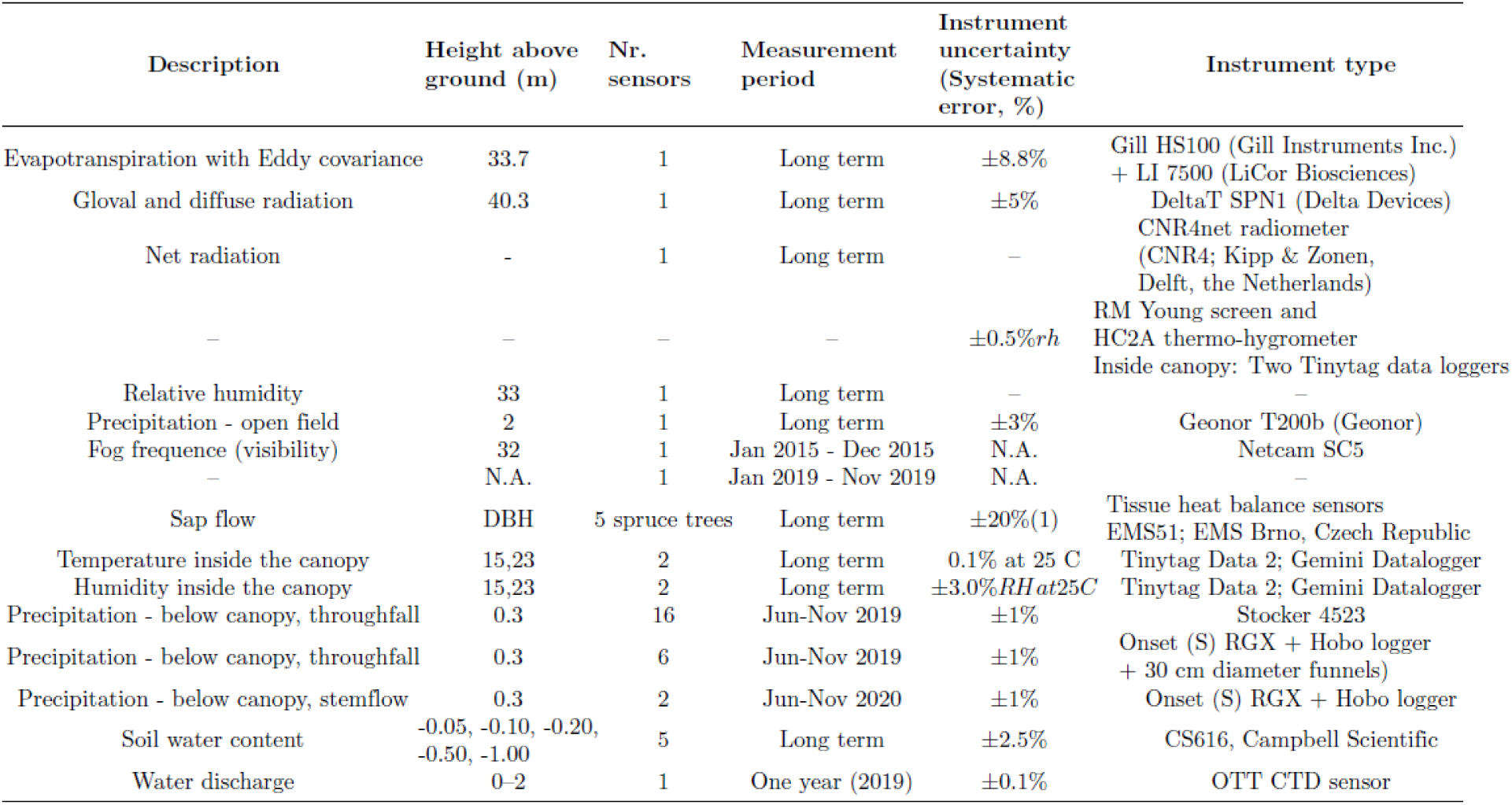
Technical description and accuracy of the measured hydrological components.

#### 2.2.1 Catchment-scale water balance

Measuring the components of the hydrological cycle at the basin scale is a typical scaling challenge (Jarvis, 1995). Although the study basin is relatively small and predominantly forested, the various methods used to assess water balance components are subject to representativeness issues. While it can be assumed that, aside from possible deep percolation, the measured water discharge quantitatively represents, in our case, basin runoff, the evapotranspiration measured using the eddy covariance technique reflects an area with a slightly different spatial footprint (see Fig. 1a, 1b). Soil water content (SWC) was measured at two locations and various depths to capture conditions under fully and partially forested cover. Sap flow, water throughfall and stemflow were measured at two representative subplots (Fig. 1c). Fog deposition and lichen-based quantifications of their water interception were performed at a single location. Despite these limitations, we believe that our setup represents the best possible compromise between instrumental cost and representativity to achieve a complete characterisation of the water cycle at the basin scale.

A 33.7 m tall tower (IT-Ren) provides data for several monitoring networks, including FluxNet (Pastorello et al., 2020), ICOS (https://www.icos-cp.eu/), and LTER (https://www.lter-europe.net/) where eddy covariance measurements are routinely performed since 1998, giving the information on evapotranspiration at a half-hour time step. The water basin surface area was measured using the local digital elevation model using ArcGIS software (ESRI, Redlands, CA, USA).

The following equation quantifies the water balance at the catchment level

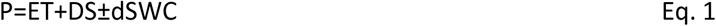

where P is Precipitation, ET is evapotranspiration, DS is water discharge from the hydrological basin, and dSWC is the variation in soil water content. Here, we quantified or estimated all the components of the water balance for the period January to December 2019.

Liquid and solid (snow) precipitation (P) was assessed by a weighing pluviometer Geonor T200x. The pluviometer was installed on a steel basement placed in a forest gap. The data were collected on a datalogger CR3000, Campbell Scientific Inc, USA (CSI). In parallel, a tipping-bucket pluviometer (Hobo, Onset, MA, USA) was also installed. Its data were collected on a CR6 CSI datalogger. The data from the two sensors were sampled every 20 seconds and averaged at 30-minute time steps.

Fog (F) was collected by a passive fog collector handcrafted at the laboratory of the University of Munich during 2015. An automatic Hobo tipping-bucket pluviometer was installed below the collector to measure intensity and timing of fog data. Fog-related meteorological conditions (vapour pressure deficit [VPD], relative humidity [RH], and diffuse/global radiation ratio) were analysed using data from 2015 and applied to 2019 using a predictive classification scheme. Days were categorised into dry, fog-only, rain-only, or mixed fog plus rain.

Evapotranspiration (ET) was assessed by the eddy covariance technique. The measuring system was composed of a 3D ultrasonic anemometer (Gill HS-50, Lymington, UK) and an enclosed-path infrared (IR) gas analyser (Li 7200 LiCor, Lincoln, NE, USA) and mounted on top of the IT-Ren tower. Air samples were taken through an insulated steel tube of 4 mm internal diameter and 0.75 m length at 0.15 m from the anemometer. The airflow rate was set to 12 L min^−1^, as provided by the flow module (7200–101, Li-Cor). All the setups followed the ICOS methodology (Rebmann et al., 2018). Raw CO_2_ and H_2_O concentration values and 3D wind speed components were measured at 20 Hz. The resulting fluxes were computed and logged by a personal computer every 30 min. and elaborated with EddyPro software (version 6.2.1, LiCor). Based on the energy balance ratio (EBR) values computed monthly (Wohlfahrt et al., 2009), we estimated the systematic errors in the eddy covariance measurements for LE+H at the Renon site to be 17.6%. Given a Bowen ratio of approximately 1 in the summer, this error can be assumed to be half of this value in LE (8.8%; Table 2) and, therefore, also in ET (Table 2). The random error of ET_EC and the other water fluxes are shown in Table 2.

Runoff (R) at the catchment scale was measured using a combination of a water stage sensor and flow velocity measurements performed with the salt dilution method. The water stage sensor was placed at the lowest spot of the catchment, just above an artificial water basin (46°35′00′′N, 11°26′02′′E, 1,675 m a.s.l.) and continuously measured the height of the water table in the stream (S in cm). Discharge (DC in L s^−1^) was calculated as

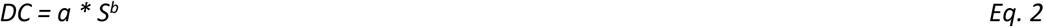

where *a* and *b* are empirical parameters according to the salt method (Day, 1977). Continuous water depth measurements were conducted in the creek with an OTT PLS 500 (OTT HydroMet Corp, VA, USA). Stream flow velocity was assessed monthly and in two more instances during 2019 after heavy precipitation events. The water discharge amount was referred to the surface of the water basin, considering its overall surface. The resulting equation describing the correlation between the stage of water (S, cm) and the discharge (D, L s^−1^) was D = 4 × 10⁻⁵ S⁴·⁵⁰⁴. The correlation coefficient R^2^ was 0.991.

The variation in water content, dSWC, was quantified using measurements from time-domain reflectometry (TDR) sensors (Campbell Scientific CS650). The sensors were placed in two locations, one in a dense forest patch (Fig. 1b) and the second in a grass patch. At both locations, they were placed at depths of 5, 10, 20, 50 and 100 cm below the litter layer. The volumetric water content (Vol Vol^−1^) variation in the first meter of depth was quantified by averaging the SWC at the different depths, linearly interpolating the values obtained and then assessing the SWC difference by subtracting the resulting water amount at the time T1 from a given water amount at the time T2, and then related to the corresponding water basin surface.

#### 2.2.2 Plot-scale water partitioning

To capture sub-plot-scale variability in forest structure and associated hydrological processes in 2019, water inputs above the canopy, the contribution of epiphytic lichens to canopy water storage and water fluxes below the canopy were measured in the studied plot.

The precipitation data used were those from the two sensors as described above. Fog occurrence was identified based on visibility <1 km, using a combination of phenocam-derived photos and public webcam imagery (www.foto-webcam.eu/webcam/ritten/) located 3 km away and 300 m below the site. The role and contribution of fog were estimated by its impact on throughfall, following the method in Hutley et al. (1997). This method was chosen due to the discontinuity of the obtained data by the fog collector during 2019. We first determined a linear regression equation between throughfall and precipitation rates for mixed precipitation and rain-only events for both plots. With this method, we obtained an estimate of predicted throughfall for a given precipitation event with rain only and with fog (mixed precipitation, Pm). Fog contribution to throughfall for mixed precipitation days was then estimated as the difference between measured throughfall and the contribution of rain to throughfall calculated as the product of precipitation and the slope of the throughfall to precipitation equation from rain-only events.

At the plot scale, evapotranspiration in both plots, old and young, was partitioned into canopy interception (I), tree transpiration (T), and soil/understory evapotranspiration. To estimate I, we used the water balance at canopy level on a daily scale during periods with rainfall-only and mixed precipitation,

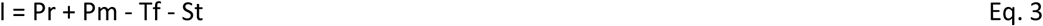

where Tf is throughfall, and St is stemflow in mm.

Tf was measured with sixteen manual rain gauges, arranged in two groups of eight in the two forest plots (Fig.1b). These pluviometers, with a 10-cm diameter orifice, were arranged in rows with a 5-m distance between each pluviometer and data were recorded almost every week. Additionally, six tipping bucket pluviometers (with a Hobo data logger; Onset) arranged 5 m apart in two groups of three continuously recorded the below-canopy precipitation at a 10-minute resolution. To increase their collection representativeness, funnels with a diameter of 30 cm were placed above the orifice of each pluviometer (Fig. 2). St was measured at one tree each at the old and young forest. Water running downwards around the tree trunk was collected into a funnel and measured with a tipping bucket pluviometer (Table 2). The measured volume of stemflow was divided by the projected crown area of the trees to convert the data into mm. Missing values owing to logger failure were added via linear correlations with automatic throughfall measurements (R² = 0.81 for the old forest patch and 0.55 for the young forest patch). The accuracy of the equipment of the measured variables and the systematic error of I in Eq. 2 are shown in Table 2. Data gaps owing to logger failure were filled by linear correlation with working gauges, as the correlations between them were high (R² > 0.87). The data for the manual gauges were missing for the last sampling interval and were added via a linear correlation with the results from the automatic gauges (R² > 0.77).

**Figure 2.**
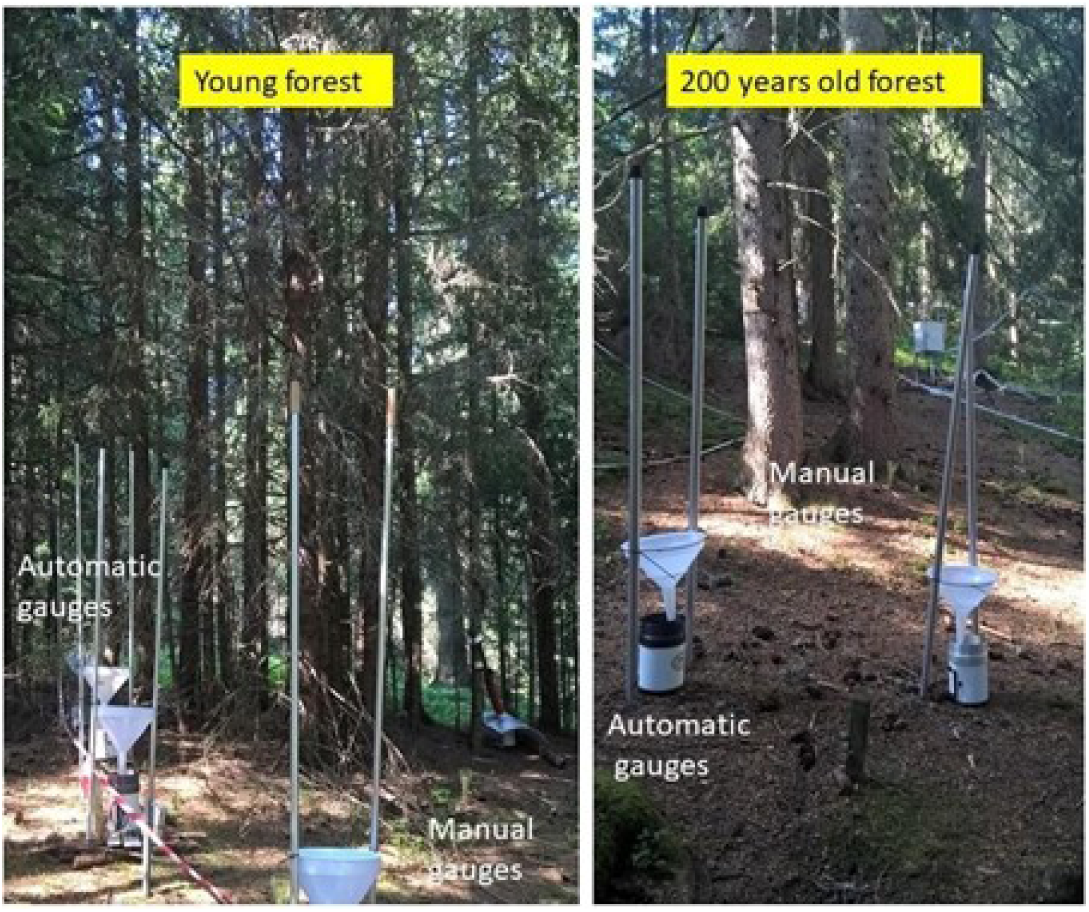
Equipment installed at ground level in both old and young forest patches to measure throughfall (automatic and manual gauges).

T was estimated by measuring the sap flow of five spruce trees with a DBH ranging from 23 cm to 57 cm. The sap flow measurements of up to 10 trees going back to 2016 showed a representative behaviour of these trees for their size classes. To minimise errors due to incoming shortwave radiation, one sensor of the tissue heat balance sensor (Table 2) was installed at the north side of the trees. The measuring system provided sap flow rates integrated for the whole sapwood depth per unit trunk circumference (kg h^−1^ cm^−1^). We logged the measured values at 10-min intervals and then scaled them to the tree level by multiplying them with stem circumference (minus the bark and phloem thickness measured during sensor installation) and integrated them to 30-min and daily sums per tree (L day^−1^) according to Kučera (2010). When a sensor was installed for more than a year, we checked whether wound reaction and/or an accumulation of resin led to a decrease in sap flow by comparing the 95^th^ percentile (P95) of 30-minute sap flow rates for 2019 and the previous years. For the two smallest trees, we found a considerable decrease in the P95 sap flow and consequently corrected the sap flow for 2019 by multiplying it by the ratio of P95 [(year of sensor installation)/P95(2019)]. Minor data gaps caused by power outages or temporary sensor malfunction were filled by using a linear correlation of the respective tree’s sap flow with eddy covariance evapotranspiration (R² = 0.57 to 0.66 at a 30-min resolution).

Tree transpiration was calculated by dividing sap flow by the projected crown area of the respective tree, thereby converting units from kg per tree to mm. The projected crown area was estimated from the mean crown radius measured in the four cardinal directions. The average tree transpiration of the two smaller trees (DBH 23 cm and 32 cm, respectively) was used to represent the young forest patch, while the average of the three larger trees (DBH 43 cm, 50 cm, and 57 cm, respectively) was used to represent the old forest patch. All the selected trees were dominant or codominant. To enable cross-scale comparison, tree sap flow measurements were scaled up to the plot level using allometric relationships and forest inventory data. These were compared against catchment-scale evapotranspiration to partition total fluxes into biophysical components. Understory and soil evapotranspiration (Esu) was estimated as the residual components of the difference between eddy covariance ET and the sum of measured transpiration and interception. The overall uncertainty in ET partitioning was derived using standard error propagation (i.e., the square root of the sum of squared component errors), ensuring consistent flux attribution across scales and enabling robust comparison of plot- and catchment-level behaviour.

The contribution of epiphytic lichens to canopy water storage was assessed from a tree that had naturally fallen in winter, with a tree height of 28 m and a DBH of 53 cm, which is representative of the old forest patch. The meteorological conditions present where the lichens were present were assessed by measuring air temperature and humidity inside the crown and the water storage capacity of the lichens at two different tree heights (Table 2).

Fresh and dry weights were determined, and water-holding capacity was calculated by difference. Values were normalised per m² of projected crown area. Lichen biomass (weight) was sampled in 3-m crown segments. In each section, all the branches were counted, and all lichens present above a single randomly selected branch were collected, together with the lichens growing on half of the main stem. In the laboratory, the lichens were first wetted until water saturation and then weighed to assess the fresh weight. They were then dried in an oven at 45 °C until a constant weight was achieved and then weighed (Sartorius Entris 2202, Göttingen, Germany) to assess their dry weight.

## 3. Results

### 3.1 Water balance at catchment level

The water balance was almost closed for the entire 2019, with a difference of 25.4 mm between the input (P = 985.6 mm ± 82.3 mm) and output ET (804.9 mm ± 70.7 mm) + DS (167.0 mm ± 12 mm) + dSWC (40 mm ± 1 mm) = 1,011.9 mm ± 103.7 mm.

The precipitation amount and seasonal pattern followed the typical pattern and amount observed in the previous 20 years of observation. The total amount of rain and snow, quantified by the Geonor T200 pluviometer, was slightly above average, with 985 mm, (the 20-year average ± standard deviation was 894±187 mm). 144 precipitation days were recorded during the year 2019 and the maximal daily rainfall was observed during the summer maximum, with 52 mm d-1 (Fig. 3a).

**Figure 3.**
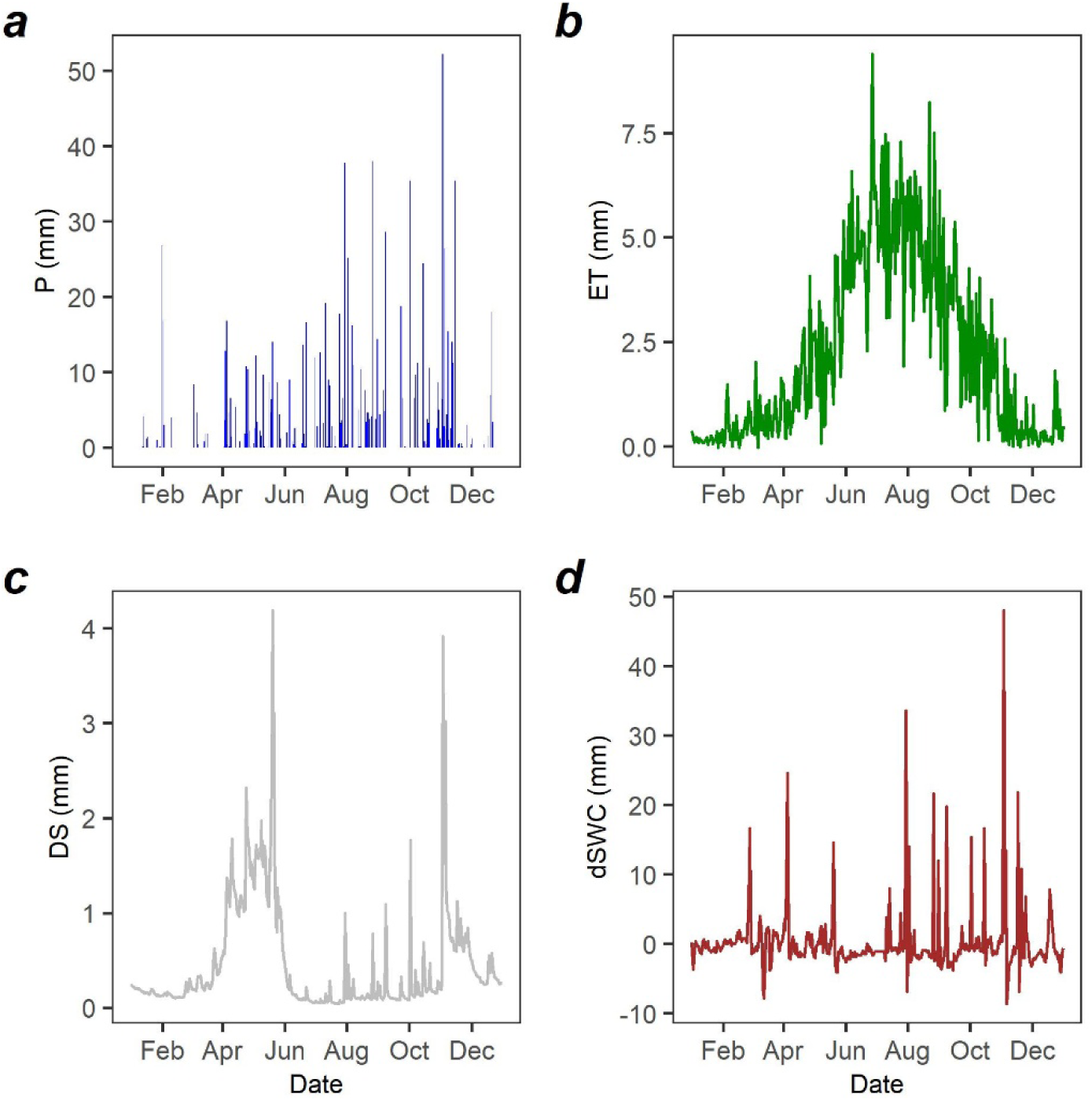
Annual pattern of the main water balance components: Top left: Precipitation, P; top right: evapotranspiration, ET; Bottom left: Water discharge, DS; bottom right: Variation in soil water content (dSWC). Measurement year: 2019.

ET values showed less variation daily. The ET peak was observed in July, with a value of emitted water vapour corresponding to 8 mm of liquid water. No prolonged soil water depletion was observed during periods of peak evapotranspiration (Fig. 3d).

The total amount of water discharged was 167 mm ± 12 mm. The resulting river flow (Figure 3c) showed a pattern with distinct peaks. One prolonged one was related to snow melting, lasting several weeks around April. In summer, we observed additional short-living peaks corresponding to intense precipitation events. However, these summer discharge peaks were considerably dampened compared to precipitation. For instance, in the precipitation event that occurred on July 30th, we recorded 37.8 mm of precipitation, while only 1 mm contributed to discharge over the 44-ha catchment, corresponding to the water discharge. Figure 3d shows that the daily difference between precipitation and water discharge during high-rainfall events can be partially attributable to the soil’s high water-retention capacity, characterised by high carbon content and low superficial bulk density (Badraghi et al., 2021). No prolonged soil water depletion was observed during periods of peak evapotranspiration (Fig. 3d).

On the annual scale (Figure 4), the balance between cumulated water input through precipitation and water losses via the sum of evapotranspiration, discharge of water from the basin and the variation in soil water content was almost perfectly closed, albeit with a possible small additional input to be added for a perfect balance, yet largely within the confidence error interval.

**Figure 4.**
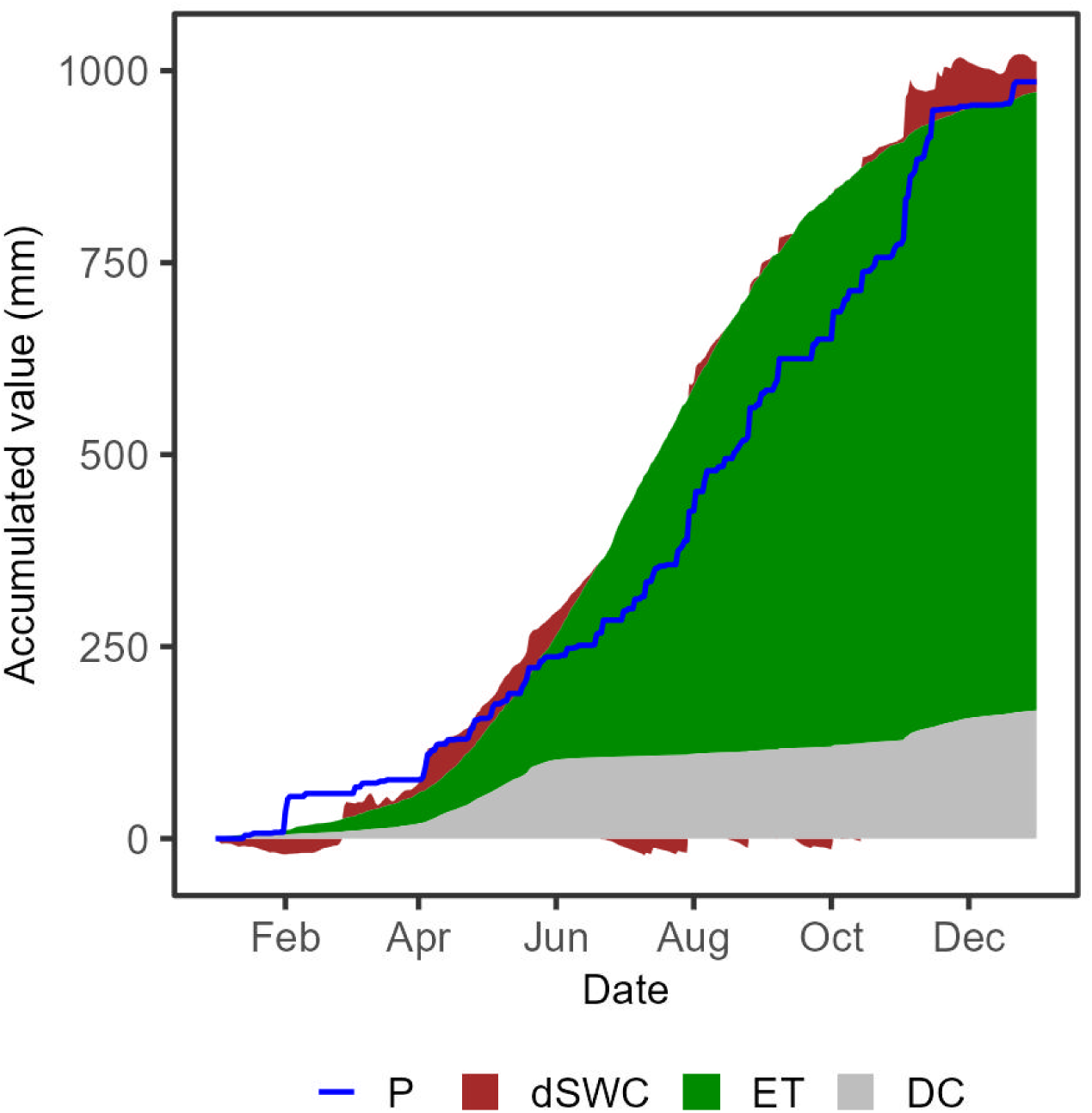
Comparison of the main water balance components: Precipitation (blue line) compared with the sum of variation in dSWC, ET and DC. Measurement year: 2019.

Overall, the outgoing fluxes were dominated by ET, representing 79.5% of the total, while DS represented 16.5%. During 2019, we measured a contribution of 4% of the left side of the water balance equation (Eq. 1) from the positive variation of dSWC, a value expected to become negligible in the long term.

### 3.2 Ecosystem evapotranspiration and tree transpiration in the young and old forest patches

The total annual evapotranspiration, estimated by the eddy covariance technique, was 808 mm. Following the same course as the previous years, the annual ET pattern was skewed, with a maximum in late summer. The large sensible heat emission dominating over latent heat in winter and spring can explain this skewness. A Bowen ratio above 1 was observed in winter and early spring, indicating that the available energy was mainly used to sustain latent heat (and ET) only in summer and autumn. Maximal daily ET was observed in summer, with a value of 9.4 mm d^−1^ (Fig. 5 a).

**Figure 5.**
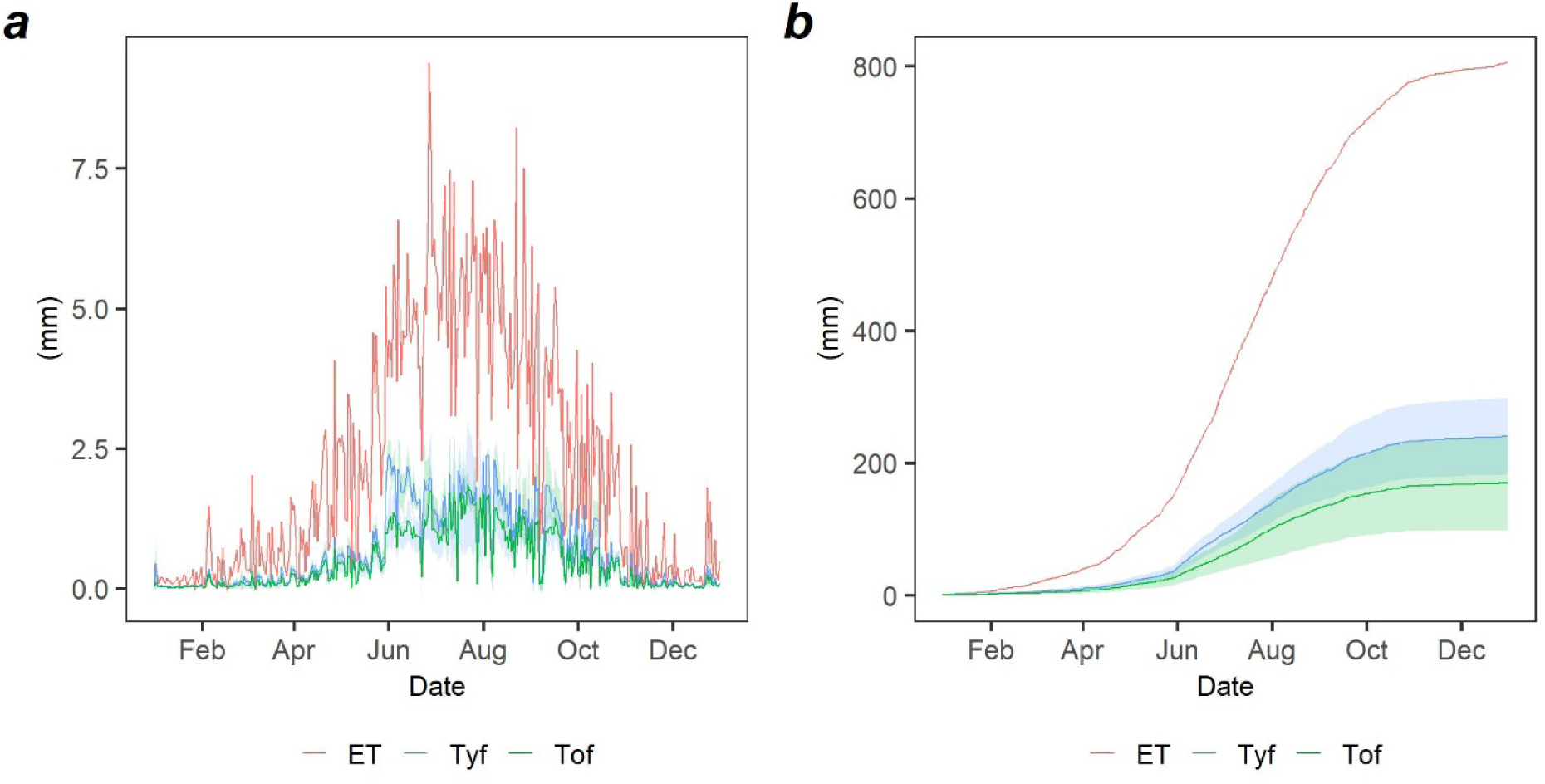
Annual pattern of evapotranspiration measured at plot scale by eddy covariance with transpiration assessed by sap flow in the young (Tyf) and old (Tof) forest patches: Figure 5a shows the daily values, Figure 5b the cumulated ones during the year 2019.

The annual pattern of ET, measured by eddy covariance, and transpiration assessed with the sap flow method measured at the plot scale, is shown in Figure 5. Transpiration in the young and old forest patches is assessed separately. We can notice that the pattern of the fluxes is consistent with ET, the R2 of the relationships being 0.85 for the young and 0.56 for the old patches. What is noteworthy is the low fraction of T from the trees as compared with total ET. On the annual scale, the transpiration observed in the young patch is 29.8% of ET, and in the old patch is 21.1%, see Figure 5b.

### 3.3 Plot-scale precipitation partitioning under contrasting meteorological conditions and the effect of forest age

#### 3.3.1 Assessment of the typical fog occurrence at the site

Long-term fog data were not available directly at the study site. Therefore, the data obtained from the webcam photos for the years 2015 and 2019 (more details in Table 3) are valuable. From January until August of 2015, the total fog frequency was 296.5 h during 72 days with fog and 37 days with mixed precipitation (109 days; 30% of days were foggy and 15% had mixed precipitation). In 2019, days with fog were frequent but there were few days with fog-only (frequent in 2015); instead, the fog was often present during days with precipitation (we counted these as days with mixed precipitation). Fog occurred for 241 h during 9 days with fog (5%) and 45 days with mixed precipitation (27%) within a measuring period of 167 days (from late spring to autumn of 2019). These results highlight the importance of days having both fog and rainfall (mixed precipitation).

**Table 3.** Days with dry conditions, fog and precipitation (rain- or snowfall) and mixed precipitation of fog plus rain-or snowfall for the whole observed period in 2015 and 2019 and the common period in both Years.

|  | 2015 all |  | 2015 common period |  | 2019 all |  | 2019 common period |  |
| --- | --- | --- | --- | --- | --- | --- | --- | --- |
| measuring period | 1.1. - 30.8.2015 |  | 25.5. - 30.8.2015 |  | 25.5. - 7.11.2019 |  | 25.5. - 30.8.2019 |  |
| time observed | 9-17 |  | 9-17 |  | 0-24 |  | 0-24 |  |
|  | days | % | days | % | days | % | days | % |
| dry | 123 | 50.6 | 40 | 41.2 | 79 | 47.3 | 46 | 47.4 |
| fog | 72 | 29.6 | 31 | 32.0 | 9 | 5.4 | 2 | 2.1 |
| rain/snow | 11 | 4.5 | 6 | 6.2 | 34 | 20.4 | 31 | 32.0 |
| mixed precipitation | 37 | 15.2 | 20 | 20.6 | 45 | 26.9 | 18 | 18.6 |
| total | 243 | 100 | 97 | 100 | 167 | 100 | 97 | 100 |

#### 3.3.2 Meteorological conditions during days with rain, fog, and mixed precipitation in 2019

Average values of radiation, relative humidity and vapour pressure deficit (VPD) during days with precipitation, fog and mixed precipitation in 2015 were calculated to predict fog occurrence in 2019 (S1, supplementary information). The ratio of total to global radiation (ratio Rg dif/Rg tot) was lower during dry periods when total Rg was high. On the contrary, diffuse Rg was less affected by rain and fog. As expected, VPD was higher and relative air humidity (RH) lower during periods with dry conditions compared to periods with fog and precipitation. Data variability was higher within the Rg dif/Rg tot ratio than within the RH and VPD.

Looking at time courses, air temperature (T) and VPD were overall lower during foggy and wet days and increased during the first dry days, but decreased again towards the end of the measuring period (Figure S2, supplementary information). Relative humidity exhibited the opposite pattern of T and VPD.

#### 3.3.3 Throughfall and canopy interception during days with precipitation and mixed precipitation

Throughfall rates relative to precipitation were higher in the young than in the old forest (Fig. 6a, b), even though the LAI in both forests was similar. Throughfall variability was higher in the manual gauges than in the automatic ones as they covered a higher small-scale variability of PAI/LAI (Fig. 6a). In addition to the automatic measurements, throughfall collected with manual gauges showed consistent patterns. Cumulative throughfall measured approximately weekly with the manual gauges in the young and old forest patches followed the same temporal evolution as the automatic measurements (Fig. 6c). When aggregated over the full observation period, cumulative throughfall derived from the manual gauges closely matched the values obtained from the automatic gauges (Fig. 6d), confirming the robustness of the observed differences between forest patches.

**Figure 6.**
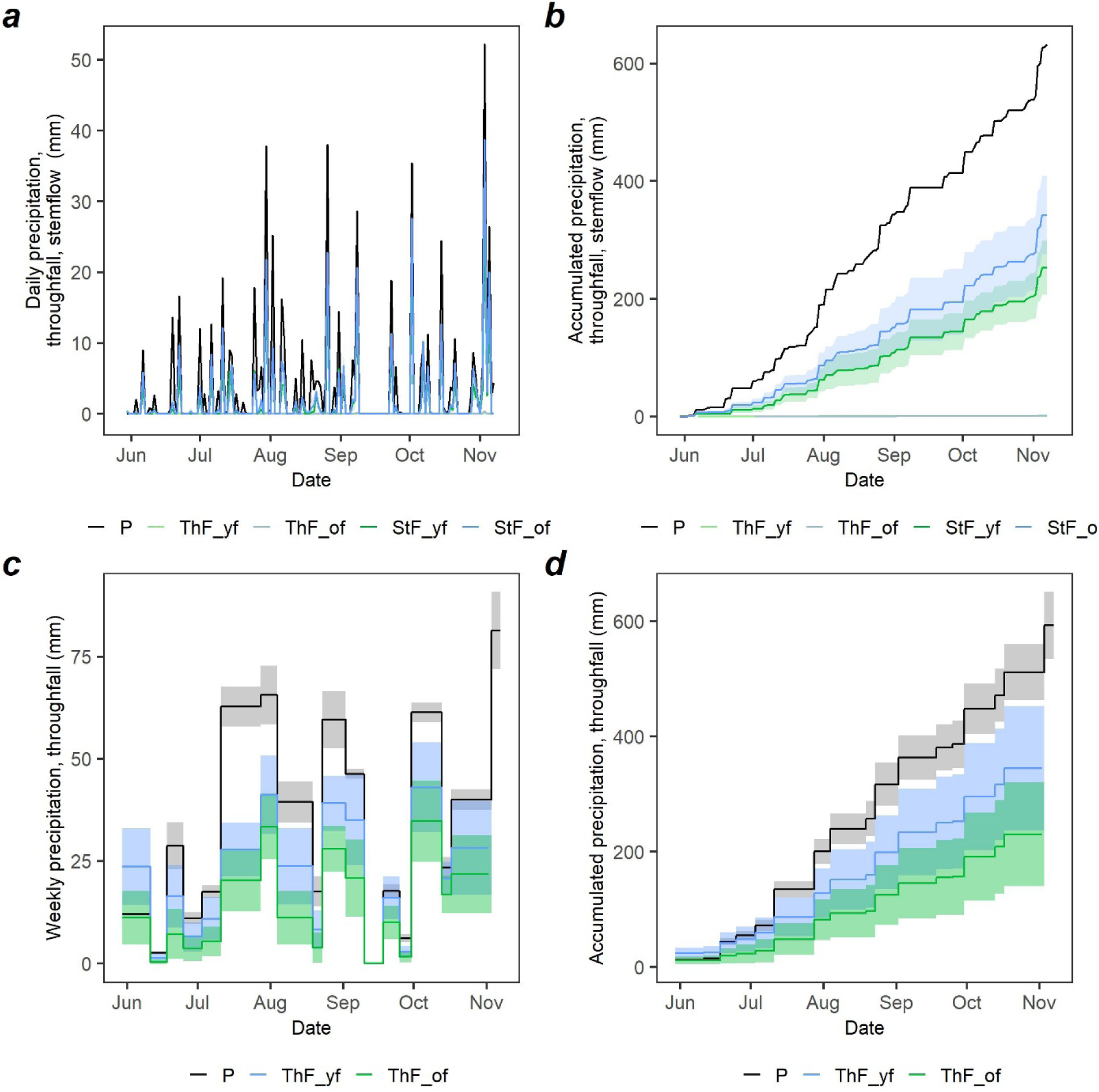
Pattern of precipitation and throughfall measured during summer and autumn of the year 2019. Figure 6a shows the pattern of precipitation and throughfall measured by the six Hobo pluviometers (three in the old forest patch, green colour, three in the young forest patch, blue colour. Figure 6b shows the cumulated values of precipitation and throughfall during the same period. Figure 6c shows the cumulated values measured approximately weekly by sixteen manual pluviometers, eight in the old and eight in the young forest patches, during the same period. Figure 6d shows the cumulated values computed using the throughfall data from the manual pluviometers.

A strong linear correlation (R² > 0.93) was found between throughfall and precipitation for both patches, and no clear increase in throughfall ratio with P. Only the last data point with the highest amount of throughfall and P was clearly above the linear regression line, indicating that the limits of the canopy’s interception capacity were reached. The correlation between old and young patch throughfall was very high and consistent across manual and automatic measurements (Figure S3, Supplementary Information), confirming that observed differences between forest patches were robust to measurement approach.

By looking at the same water components, split into approximately monthly periods (adjusted to the sampling periods of the manual throughfall gauges), it can be noticed that P increased from June to October, while ET was high until September, but decreased strongly afterwards (Table S4, Supplementary information). The Tf/P ratio was higher in autumn (September and October) than in summer, except for June in the old forest; correspondingly, the interception rate (I/P) was lower. This should be associated with higher P, but also with more mixed precipitation days observed in autumn. Additionally, stemflow, which was similar for both forest plots, increased from June to October but was overall too low to play a major role in the water balance.

By looking at throughfall rates at a daily resolution and separating by precipitation type, we found that throughfall was much higher during days with mixed fog and rain precipitation (Figure 7; Table 4), conversely interception rates (I/P) were lower during rain-only days. This surplus in Tf (Tf/P = 0.28 for the young forest and Tf/P = 0.27 for the old forest), was attributed to fog.

**Figure 7.**
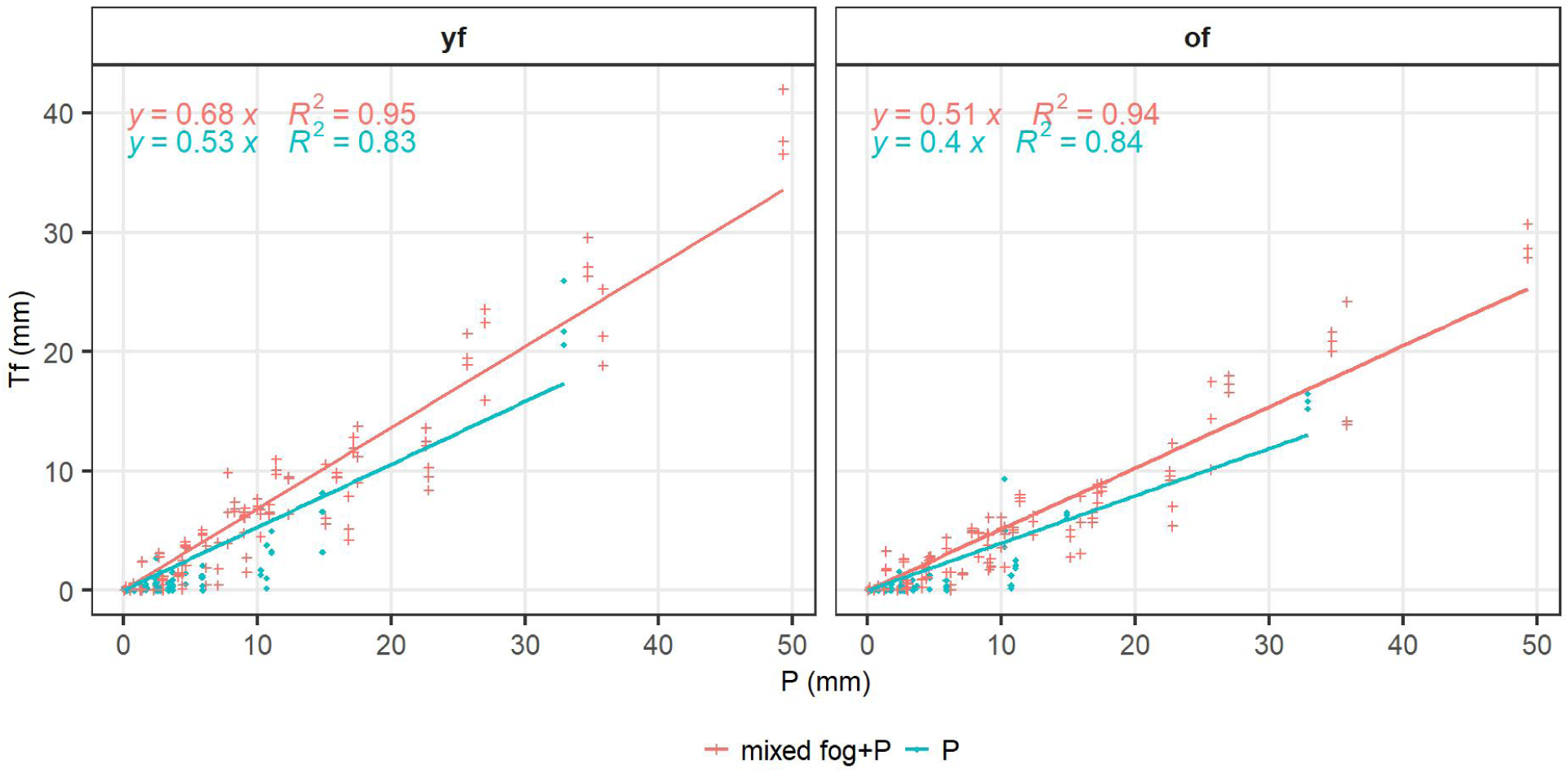
Throughfall versus precipitation during mixed precipitation (mixed fog+P) and rain-only (P) events in the young (yf) and old (of) forest patches.

**Table 4.** Estimated fog contribution to throughfall in mixed precipitation events in the young and old forest patch (mean ± standard deviation for absolute amounts). Fog contribution was estimated daily for all single throughfall gauges and negative estimations were set to zero, thus the sum of estimated fog and rain contribution was higher than measured throughfall.

| Days with mixed precipitation | Young forest patch | Old forest patch |
| --- | --- | --- |
| P measured (mm, see Table 3) | 460 ± 35 | 460 ± 35 |
| Total Tf measured (mm) | 292 ± 26 | 216 ± 11 |
| Tf estimated from rain only events (mm) | 243 ± 7 | 184 ± 8 |
| Fog contribution in mixed events (mm) | 70 ± 15 | 53 ± 5 |
| measured Tf/P (%) | 64 | 47 |
| estimated rain-only Tf/P (%) | 53 | 40 |
| estimated fog Tf/P (%) | 15 | 12 |
| Rain contribution to Tf (%) | 83 | 84 |
| Fog contribution to Tf (%) | 24 | 24 |

## 4. Discussion

### 4.1 The water balance and evapotranspiration partitioning across scales

The annual water balance at the Renon catchment was almost closed in 2019, with a small residual of 25.4 mm between precipitation (P = 985.6 mm) and the sum of evapotranspiration (ET = 804.9 mm), discharge (DS = 167.0 mm), and the net increase in soil water storage (dSWC = 40 mm) (Fig. 3; 4). In relative terms, ET accounted for 81.7% of P and DS for 16.9% of P (dSWC 4.1% of P), confirming a strongly ET-dominated catchment water balance in 2019 (Fig.4).

This dominance of evapotranspiration was consistent with the event-scale discharge/runoff response, which remained small even for larger rainfall events. For example, in Fig. 3, a rainfall event of 52 mm produced only 3.5 mm of discharge, and on 30 July 2019, a precipitation depth of 37.8 mm generated only 1.0 mm of discharge (44 ha catchment). These responses indicate that annual discharge remained modest relative to atmospheric losses, showing a substantial buffering by subsurface storage and a limited fraction of precipitation translating into quick flow under the observed conditions at the catchment scale.

At the ecosystem/stand scale, eddy-covariance evapotranspiration peaked at 9.4 mm d⁻¹ and remained systematically higher than sap-flow-derived transpiration in both forest types (Fig. 5). The seasonal decrease is consistent with energy partitioning constraints, as elevated Bowen ratios outside the main summer period indicate a larger sensible-heat share (Fig. 3). Rainfall events occurred regularly across the growing season with only short dry spells, and soil moisture dynamics shown in the catchment monitoring indicate no extended drought period during the main measurement window (Fig. 3). Neither ET nor T showed a clear event-scale response to precipitation inputs (comparison with Fig. 3), indicating that atmospheric demand and canopy processes dominated short-term evaporative dynamics during the measurement period.

The relationship between transpiration and ET was coherent in time (Fig.5), indicating a shared meteorological control but different partitioning of ET components between stands. Sap-flow-derived transpiration (T) represented only 29.8% of ET in the young and 21.1% of ET in the old forest patch. Low T/ET ratios have been reported across conifer forests with a broad range (15–75%) depending on site conditions, stand structure, and methods (Cienciala et al., 1992; Grelle et al., 1997; Köstner, 1998; Lagergren et al., 2008; Matyssek et al., 2009; Brito et al., 2015; Soubie et al., 2016). At Renon, the low T/ET values implied that 70–79% of ET ((ET − T)/ET) was attributable to non-transpiration components (dominated by wet-canopy evaporation and evaporation from intercepted water, plus understory/soil evaporation). The magnitude of “missing” ET relative to transpiration is consistent with interception theory and empirical studies: forests commonly intercept 10–30% of rainfall, but can reach up to 50% depending on canopy structure and storm characteristics (van Dijk et al., 2015). In our dataset, stemflow was negligible, consistent with conifer-forest studies where stemflow typically contributes little to net rainfall (Dohnal et al., 2014).

For cross-scale interpretation, the stand-scale precipitation partitioning implies substantial wet-canopy evaporation potential during precipitation days and mixed rain–fog conditions. During days classified as mixed precipitation, gross precipitation was P = 460 ± 35 mm, while measured throughfall amounted to 292 ± 26 mm (young) and 216 ± 11 mm (old), with stemflow of only 1.0 mm (young) and 0.8 mm (old) (Table S5). The implied interception losses were 167 ± 36 mm (young; 36% of P) and 242 ± 29 mm (old; 53% of P), demonstrating a marked structural control on interception.

These values are plausible for a spruce mountain context: Dohnal et al. (2014) report interception losses of 33–36% for Norway spruce for comparable seasonal windows and show strong sensitivity to stand structure. The fact that our old forest reaches the upper end (53% of P in mixed events) supports the interpretation that denser/structurally more complex canopies sustain longer wetting–drying cycles and higher wet-canopy evaporation, which helps reconcile high EC-derived ET with comparatively modest sap-flow transpiration.

Differences between forest patches further highlight the role of stand structure in regulating transpiration at the plot scale. Transpiration per unit ground area was higher in the young than in the old forest patch, despite larger individual trees in the old patch (Fig. 5). Higher stem density and smaller projected crown areas in the young forest (Table 1) compensated for lower sap flow rates of individual trees, resulting in greater stand-level transpiration. While transpiration in both forest patches showed coherent seasonal responses to atmospheric conditions (Fig. 5), the slope of the relationship between transpiration and ET was higher in the young forest, reflecting differences in flux partitioning rather than contrasting climatic sensitivity.

At the Renon site, two mechanisms explain the relatively small contribution of transpiration to total ET. First, canopy interception evaporation represents a major non-physiological evaporative pathway, particularly in the old forest patch, where interception to precipitation ratios are high (Fig. 6; Table 4). Intercepted water is rapidly returned to the atmosphere without stomatal control, thereby reducing the relative share of transpiration even during periods of active vegetation. Second, understory evapotranspiration and soil evaporation contribute substantially to ecosystem-scale ET, as indicated by the residual between eddy covariance-derived ET and the sum of measured transpiration and estimated interception (Table S5). Comparable contributions of understory and soil fluxes have been reported in other conifer forests, where their magnitude depends on ground vegetation cover, light penetration, and soil moisture dynamics (Köstner, 1998). In addition, at the Renon site, the irregularity in the forest cover may have contributed to this result.

This cross-scale evidence is consistent with the broader notion of subalpine, forested “water towers” in which a large fraction of incoming water returns to the atmosphere and only a smaller proportion is exported as runoff (Messerli et al., 2004; Viviroli et al., 2007), and comparable or higher ET fractions have been reported for coniferous mountain forests in the European Alps and other temperate regions under warm summers and high canopy cover (Grelle et al., 1997; Soubie et al., 2016; Mastrotheodoros et al., 2020). Uncertainty in absolute transpiration estimates remains relevant when interpreting low T/ET ratios, as sap flow is sensitive to sensor performance, wound responses, radial variability of sap flux, and scaling assumptions (Wilson et al., 2001; Schlesinger and Jasechko, 2014; Peters et al., 2018; Flo et al., 2019). In this study, consistency checks include the coherence between sap-flow-derived transpiration and eddy-covariance ET (Fig. 5) and the applied correction approach (sap flow correction), which reduce but do not eliminate uncertainty in absolute magnitudes.

### 4.2 Forest age, structure, and interception dominance

The plot-scale partitioning results demonstrate that forest age exerts a strong control on how precipitation is redistributed within the canopy and partitioned into throughfall, interception, and transpiration. Despite similar leaf area indices (LAI ≈ 4.8) in both forest patches, throughfall and transpiration were consistently higher in the young forest, whereas interception was markedly higher in the old forest. Over the observation period, interception accounted for 54 % of total precipitation in the old forest, compared to 33 % in the young forest, indicating that canopy structure and surface water storage capacity associated with stand development are more important than leaf area alone in governing interception efficiency.

Event-based description provides additional insight into evapotranspiration controls. Dry days contributed disproportionately to total ET, whereas days characterised by mixed precipitation and fog exhibited suppressed ET (Table S5). This suppression is consistent with reduced vapour pressure deficit and lower radiation under humid and cloudy conditions (Fig. S1; Fig. S2) and aligns with observations from cloud-affected forest ecosystems where fog presence dampens evaporative losses (García-Santos et al., 2009; García-Santos et al., 2012). At the Renon site, however, mixed precipitation days occurred more frequently later in the growing season (Table 3), so seasonal co-variation between fog occurrence, phenology, and energy limitation must be considered. Consequently, fog is best interpreted here as a systematic modifier of micrometeorological conditions, rather than as a dominant driver of annual evapotranspiration.

As intercepted water is returned directly to the atmosphere without passing through the soil or plant stomata, high interception fractions in the old forest reduce the proportion of precipitation available for soil recharge and transpiration, while enhancing non-stomatal evaporation. This shortcut in the water cycle provides a mechanistic explanation for the lower transpiration-to-ET ratios observed in the old forest (T/ET ≈ 22 %) compared to the young forest (T/ET ≈ 31 %), as discussed in Section 4.1.

Higher throughfall in the young forest patch resulted in a greater fraction of precipitation reaching the forest floor, supporting soil moisture recharge, understory evapotranspiration, and potential runoff generation. Stand-level transpiration was also higher in the young forest, despite lower sap flow rates of individual trees, due to higher stem density and smaller projected crown areas. When scaled to the plot level, these structural characteristics resulted in higher cumulative transpiration in the young forest. In addition, the residual component of evapotranspiration attributed to soil and understory evaporation (calculated as ET − T − I) was larger in the young forest, consistent with lower interception losses and greater water availability below the canopy.

Although both forest patches experienced identical meteorological forcing, their contrasting partitioning of precipitation underscores the role of structural legacies in shaping hydrological behaviour. In the old forest, a larger fraction of incoming precipitation is rapidly cycled between canopy and atmosphere via interception evaporation, whereas in the young forest a greater proportion of water is routed through the soil–plant continuum. At the catchment scale, such age-related differences in interception and throughfall have the potential to influence the timing and magnitude of soil water recharge and streamflow response, even when total water losses are dominated by evapotranspiration. These results demonstrate that forest age and structure must be explicitly accounted for when upscaling plot-scale observations or assessing hydrological consequences of forest management and succession.

### 4.3 Epiphytes as modifiers of canopy water storage and energy balance

Despite similar LAI in both forest patches, precipitation partitioning differed strongly between patches. Throughfall was systematically higher in the young forest, whereas interception/storage was markedly higher in the old forest during the same observation window (Fig. 6; Table S5). This divergence indicates that leaf area alone is insufficient to explain interception dominance and points to age-related differences in canopy structure and surface storage capacity. This is consistent with interception theory, where canopy roughness, surface wettability, storage capacity, and wet-canopy evaporation regulate net rainfall and interception loss (Savenije, 2004; van Dijk et al., 2015).

A plausible mechanism for the elevated interception/storage in the old forest patch is the presence of epiphytic lichens and other non-vascular epiphytes that expand the wettable surface area and provide an additional water-holding compartment on branches and stems. Epiphytes are known to increase canopy water storage and prolong wet-canopy conditions, thereby enhancing evaporation of intercepted water and altering energy partitioning (Pypker et al., 2006; Sillett and Van Pelt, 2007; Porada et al., 2018). Importantly, epiphyte effects can be hydrologically relevant even when not dominant in the annual catchment balance. They primarily modify how precipitation is redistributed within the canopy and how long canopy surfaces remain wet, which directly controls the magnitude of interception loss (van Dijk et al., 2015).

Our dataset supports the feasibility of this pathway in two independent ways. First, within-canopy micrometeorology indicates a persistently humid crown environment that would prolong epiphyte hydration and slow drying. Mean RH was higher and mean VPD lower inside the canopy (especially at 15 m) than outside (Table S6). These conditions are consistent with longer wet-surface residence times and thus greater opportunity for evaporation from stored canopy water rather than drainage as throughfall (Holwerda et al., 2012; van Dijk et al., 2015). Second, we quantified lichen water-holding capacity on a representative old-forest tree (DBH 53 cm; 28 m height) using fresh–dry mass differences and crown-segment sampling. While this sampling design does not allow a stand-level epiphyte inventory, it provides site-specific evidence that the old canopy contains a measurable lichen storage pool and therefore can plausibly exhibit larger effective storage capacity than the young patch. This interpretation is consistent with the observed water partitioning contrast (Table S5).

The epiphyte mechanism also complements the evapotranspiration partitioning inferred in Sections 4.1–4.2. At Renon, sap-flow-derived transpiration accounted for only ∼30% (young) and ∼21% (old) of ecosystem ET, implying a large non-transpiration component. Interception evaporation is a primary process for this residual, particularly in the old forest where I/P is highest (Table S5). This is consistent with observations that wet-canopy evaporation and interception can represent a substantial fraction of forest ET, especially in humid or frequently wetted canopies (Holwerda et al., 2012; van Dijk et al., 2015). Therefore, epiphytes are best interpreted here not as an additional water input source, but as a structural modifier that increases canopy storage and wet-surface evaporation potential, strengthening interception dominance in the old forest patch.

Limitations are acknowledged explicitly: We refrain from scaling lichen storage to a stand-level storage term because epiphytes were quantified on a single representative tree rather than across the stand. Consequently, we treat epiphytes as a mechanistic explanation supported by (i) robust age-related partitioning differences at similar LAI, (ii) within-canopy humidity and VPD conditions, and (iii) direct lichen water-holding measurements, but not as a fully parameterized canopy-storage model component.

### 4.4 Fog as a secondary but systematic modifier of precipitation partitioning

In temperate mountain forests, fog and cloud water are difficult to quantify as a separate hydrological input because fog frequently co-occurs with rainfall, varies with exposure and turbulence, and is sensitive to measurement approach (Katata, 2014). At Renon, fog-only throughfall was not captured, and therefore, we do not estimate a stand-level fog-only input. Instead, we adopt a conservative strategy. We evaluate fog as a secondary but systematic modifier of precipitation partitioning by focusing on mixed fog–rain events, where fog presence is detectable (Table 3) and its influence can be inferred from the throughfall–precipitation relationship (Hutley et al., 1997).

Our results show that mixed precipitation was common at Renon during the 2019 observation window. Fog occurred mainly in combination with precipitation (45 mixed days versus 9 fog-only days; Table 3). Mixed events exhibit a repeatable throughfall surplus relative to rain-only expectations. Using regressions derived from rain-only events (Fig. 7), we estimate an additional throughfall component attributed to fog of 70 ± 15 mm in the young forest and 53 ± 5 mm in the old forest over the mixed-event set (Table 4). Expressed as fractions, fog contributes ∼12–15% of incident precipitation during mixed periods but ∼24% of throughfall in both stands (Table 4). Fog is not an occasional anomaly but a recurrent modifier that measurably increases below-canopy inputs when it co-occurs with rainfall.

We did not capture throughfall during days with fog only. In subtropical cloud forests with a high frequency of fog, fog-only contributed only 6% of throughfall (García-Santos, 2007). This contrasts with the Japanese Alps, Mt. Tateyama, where Uehara and Kume (2012) measured up to 35% of fog-only contribution in throughfall, attributed to the high wind velocity and humidity in the Japanese Alps. The lower contribution in the Alpine forest may be explained by differences in air temperature and altitude. This result is in line with our hypothesis that fog might have a minimal impact on the annual and seasonal water budget.

The direction and magnitude of this effect are consistent with studies showing that cloud and fog deposition can increase net precipitation and throughfall relative to open rainfall, particularly under frequent cloud immersion and in conifer canopies that efficiently scavenge small droplets (Köhler et al., 2014; Yamaguchi et al., 2015). However, the Renon case also shows why fog should be treated as secondary in annual closure. The catchment water balance is nearly closed without an explicit fog term (Section 4.1), and fog-only inputs were not measured directly. Our inference is therefore intentionally limited to what the data support. Fog modifies partitioning primarily during mixed events and is expressed as a throughfall surplus.

Fog can also influence partitioning indirectly via microclimate. Foggy and mixed days are associated with lower VPD and higher RH (Figs. S1–S2; Table S6), which suppresses drying and can reduce wet-canopy evaporation rates per unit time while extending wetness duration. This provides a mechanistic link to event-class stratification, showing that different classes exhibit distinct partitioning behaviour (Table S5). It also supports the interpretation that fog affects not only how much water reaches the ground during mixed events, but also how canopy storage and evaporation operate under humid conditions. This aligns with process syntheses emphasizing that fog and cloud deposition interact strongly with canopy turbulence, wetness duration, and evaporative exchange, and that uncertainties are substantial across methods and sites (Katata, 2014).

### 4.5 Implications for ecohydrological regulation in subalpine forests

The Renon catchment exhibits a strongly ET-dominated annual water balance (Section 4.1). This is consistent with the “water tower” concept in which mountain forests recycle a large fraction of precipitation to the atmosphere and export a smaller fraction as runoff (Messerli et al., 2004; Viviroli et al., 2007; Beniston et al., 2011). Within this catchment-scale constraint, our patch-scale results demonstrate that forest age and associated canopy traits control how precipitation is partitioned at the canopy level. This has implications for below-canopy water availability, wet-canopy evaporation, and the pathways by which atmospheric inputs become soil water and streamflow.

Three implications follow directly from our measurements. First, stand development can shift the balance between canopy–atmosphere recycling and soil–plant routing. During mixed precipitation conditions, the old forest retained a much larger fraction of incident precipitation as interception/storage than the young forest (Table S5), while stemflow remained negligible in both stands. Because interception evaporation returns water directly to the atmosphere without passing through the soil, higher interception in older forest patches implies less water reaching the forest floor and potentially reduced effective recharge at the patch scale, even if the catchment remains buffered overall (van Dijk et al., 2015).

Second, fog modifies partitioning even when it is not a dominant annual input. Mixed fog–rain conditions increased throughfall systematically (Table 4; Fig. 7). This supports the interpretation that fog influences below-canopy inputs primarily through event-scale modification rather than as a separable annual flux. This has relevance under climate change because changes in cloud immersion frequency, seasonal timing, or cloud base height could shift the prevalence of mixed events and thus the frequency of throughfall enhancement and humid-canopy conditions, even if total precipitation remains similar (Gobiet et al., 2014; Katata, 2014).

Third, epiphytes likely amplify age effects by increasing canopy storage capacity and wetness duration. Our lichen storage measurements provide direct site evidence for an additional canopy storage component in the old forest patch, and the within-canopy microclimate supports longer wet-canopy persistence (Table S6). Together with the robust difference in interception dominance between patches at similar LAI (Table S5), this supports a mechanistic interpretation consistent with epiphyte interception literature. Epiphytes increase water storage and prolong canopy wetness, which enhances interception loss and influences energy partitioning (Pypker et al., 2006; Sillett and Van Pelt, 2007; Porada et al., 2018). This pathway is particularly relevant in humid, frequently wetted canopies, where wet-canopy evaporation can be a large share of ET (Holwerda et al., 2012; van Dijk et al., 2015).

Limitations and robustness should be stated clearly. Fog-only inputs are not quantified because fog-only throughfall was not captured. Accordingly, we restrict claims to mixed-event modifications inferred from throughfall–precipitation relationships and fog occurrence classification (Hutley et al., 1997). Epiphyte storage was measured on a representative tree rather than mapped across the stand. Thus, epiphytes are treated as a supported mechanism rather than a stand-parameterized storage term. The transfer of findings across scales can induce uncertainty, particularly if the stand characteristics vary across spatial scales. Nevertheless, the age signal is robust because it is replicated across manual and automatic throughfall networks and consistent across temporal aggregations (Fig. 6; Fig. S3).

## Conclusion

This study links catchment-scale monitoring with tree-scale measurements to quantify the hydrological balance of a subalpine conifer catchment and to explain how forest structure and fog occurrence shape precipitation partitioning. At the catchment scale, the annual water balance was close to closure and indicates that evapotranspiration is the dominant flux returning incoming precipitation to the atmosphere, whereas discharge represents a smaller fraction exported from the catchment. At the forest patch scale, precipitation partitioning differed systematically between old and young formations: throughfall was consistently lower in the old forest than in the young forest despite broadly similar LAI, implying that interception capacity is controlled not only by leaf area but also by structural and surface properties that alter canopy storage and wet-canopy evaporation. The presence of abundant epiphytic lichens in the old forest provides a coherent mechanism for this difference by increasing canopy water storage and prolonging wet-canopy conditions, thereby reducing the fraction of precipitation reaching the forest floor as throughfall.

Fog emerged as a secondary but systematic driver of partitioning. Although fog-only inputs could not be isolated as an independent water-balance term in this temperate mountain setting, mixed fog–rain events produced a detectable and recurring modification of patch-scale partitioning: fog occurrence can increase effective below-canopy inputs through canopy droplet capture and drip, while simultaneously modifying interception losses by sustaining high humidity and low vapor pressure deficit that slow canopy drying. These effects do not need to dominate annual precipitation totals to be hydrologically relevant; by recurring under specific seasonal and synoptic conditions, fog can systematically shift event-scale throughfall, canopy storage dynamics, and the timing of water transfer through the canopy. Overall, our results show that linking scales is essential to interpret ecohydrological functioning in subalpine conifer catchments, where forest age and epiphyte-mediated canopy properties interact with fog occurrence to shape the partitioning of precipitation into throughfall, interception, evapotranspiration, and runoff export.

## Acknowledgements

The authors thank the University of Klagenfurt for the sabbatical period of Assoc. Prof. Dr Garcia-Santos. MSc. Thomas Velina from the University of Klagenfurt for his contribution in converting the phenological photos to visibility data, Dr Gert Wolf from the University of Klagenfurt for assisting us with data analysis, Dr Jay Frentress, Dr Michael Engel and Prof. Francesco Comiti, all four from the University of Bozen/Bolzano, for providing water stage and water discharge information.

